# Community composition, and not species richness of microbes, influences decomposer functional diversity in soil

**DOI:** 10.1101/2023.04.13.536737

**Authors:** Shamik Roy, Jalmesh Karapurkar, Pronoy Baidya, M. Jose, Sumanta Bagchi

## Abstract

Diversity-function relationships are well known for producers, and these are also known to be influenced by consumers at higher trophic levels. However, these are not well known for microbial decomposers and decomposition processes in soil. Further, it also remains unknown whether and how consumers such as large mammalian herbivores, who are a major feature across more than one-third of the world’s terrestrial realm, influence soil microbial decomposer diversity-function relationships.
We used a14-year old long-term herbivore-exclusion experiment to answer two questions: (a) whether microbial functions vary with microbial diversity (both species richness and composition), and (b) whether herbivores alter the diversity-function relationships among microbial decomposers. We measured functional richness, and functional dispersion from the utilization profiles of 30 metabolic substrates. Alongside, we also measured species richness and species composition of soil microbes.
We analysed soils from n=10 paired grazed-and-fenced plots, and sampled them three times during the growth season of 2019. Data were from 60 16S rDNA gene amplicon sequences with 7.6 million reads covering 1937 operational taxonomic units (OTU) across 47 phyla (of which 924 OTUs were identifiable to genus level), and 1800 catabolic profiles.
Functional diversity in soil was positively related to microbial community composition, but not to species richness. This parallels the diversity-function relationships between producers and production; it also challenges the prevailing notion of functional redundancy in hyper-diverse soil microbial communities. Since certain combinations of species can outperform others, global change factors which can alter microbial communities may also impact decomposition processes and services. But this coupling between diversity and functions was unaffected by experimental herbivore-exclusion, indicating resilience and resistance among decomposers.
Structural equation models suggested that the strength of this relationship is favoured by availability of soil moisture. These also showed microbial functions varied more strongly with temporal variables (e.g., seasonality) than with spatial variables (e.g., edaphic factors such as soil texture and pH). Although interpretations can be constrained by which and how many functions are investigated, the relationship between diversity and functions was generalizable and robust once 16 or more functions were quantified.
While ecosystem functions and services derived from microbial decomposers have intrinsic resilience and resistance, they respond strongly to variability in water availability. Decomposition in grazing ecosystems may be particularly susceptible to how microbial community composition responds to rising precipitation variability under ongoing and projected climate change.

**Highlights:**

- Diversity-function relationships for microbial decomposers remain uncertain.
- Microbial diversity is positively related to functions
- This relationship is robust to herbivore-exclusion.
- Microbial diversity-function relationship can be susceptible to altered precipitation

## Introduction

Species diversity begets ecological functions, and high taxonomic diversity promotes functional diversity as well. Oppositely, depending on ecological resilience and resistance, loss of diversity can deplete functions. The empirical evidence for such diversity-function relationships mostly come from producers and their production (Schulze and Mooney, 1993; Loreau et al., 2001; Hooper et al., 2005; Allison and Martiny, 2008; Reich et al., 2012; Tilman et al., 2014). These diversity-function relationships among producers are also known to be modulated by consumers at higher trophic levels (Olff and Ritchie, 1998; Duffy, 2002; Duffy et al., 2007; Cardinale et al., 2011). It is pertinent to ask whether the patterns and processes seen in producers can be extrapolated to decomposers and to decomposition processes. But the answer seems to be contentious as both ‘yes’ and ‘no’ have been reported by studies examining diversity-function relationships for decomposers (Delgado-Baquerizo et al., 2016a; Louca et al., 2016; Wagg et al., 2019; Chen et al., 2022). Generally, experimentally assembled communities in lab conditions tend to show positive diversity-function relationships in decomposers (Delgado-Baquerizo et al., 2016a; Wagg et al., 2019). In contrast, when natural variation in species richness in ecosystems is weighed against functions, studies tend to find no relationship and this is attributed to functional redundancy in decomposers (Louca et al., 2016; Chen et al., 2022). Such contradictions could be due to few species in experimental studies (c. 10^1^ to 10^2^ species) (Delgado-Baquerizo et al., 2016a; Wagg et al., 2019) and many species in observational studies (c. 10^3^ species) (Louca et al., 2016; Chen et al., 2022). However, there are indications that microbial community composition—not their species richness, per se—influences decomposition functions in natural communities (Fierer et al., 2013; Delgado-Baquerizo et al., 2017; Galand et al., 2018). Since this challenges the long-standing assumption of functional redundancy in hyper-diverse communities where multiple taxa could perform the same functions, it is pertinent to ask whether microbial decomposition process in soil is related (i.e., coupled) to community composition. However, much of the available evidence comes from metagenomic predictions of putative functions, rather than from quantification of the functions themselves (Fierer et al., 2013; Galand et al., 2018). Evaluating the relationship against actual functions is critical for knowing whether diversity can buffer and ameliorate the effect of stressors that act on decomposition functions and services.

Decomposition processes run by microbes do not occur in isolation of other trophic levels. Large mammalian herbivores as consumers of plant production can simultaneously alter both the quantity and quality of biomass-input to soil. Grazing can influence microbial community composition and regulate taxonomic profiles of the decomposer community in soil (Bardgett et al., 2001; Ylänne et al., 2021). Simultaneously, grazing can also influence functional attributes of soil microbes (Bagchi et al., 2017; Roy and Bagchi, 2022; Roy et al., 2023). But it remains poorly understood whether grazer influence on microbial functions in soil is mediated through their effects on microbial community composition, or not. In addition to grazers, underlying edaphic conditions such as moisture availability (Bagchi et al., 2017), soil pH (Fierer et al., 2006; Delgado-Baquerizo et al., 2016b), and texture (Girvan et al., 2003) can also affect microbial decomposers. Moreover, soil microbial communities and functions can be influenced by seasonality as they undergo a high degree of temporal change as well (Frossard et al., 2012; Bagchi et al., 2017; Roy et al., 2023). The role of large mammalian herbivores in different abiotic conditions in mediating the inter-dependency between the taxonomic and functional aspects of soil microbial diversity can have broad implications for local, regional, and global biogeochemical cycles (Bardgett and Wardle, 2003; Bagchi et al., 2017; Naidu et al., 2022; Roy and Bagchi, 2022; Roy et al., 2023).

Here we assess microbial diversity-function relationship by measuring the variation in microbial diversity and functions at regular intervals through the growth season in the Trans-Himalayan grazing ecosystem of northern India. Specifically, we asked whether: (1) microbial diversity (both species richness and composition) and functions vary with grazing and across the growth season (early-, mid-, and late-season); (2) microbial community composition influences decomposition functions; (3) herbivores alter the relationship between functions and species composition that can be otherwise mediated by edaphic conditions and by seasonality.

## Methods

### Study site and sampling

We used a long-term replicated herbivore-exclusion experiment started in 2005-06 in Spiti region of Himachal Pradesh in northern India (32° N, 78° E) (Bagchi and Ritchie, 2010a). We set up replicated herbivore exclosures, each 10×10 m^2^, with adjacent paired control grazed plots. Spiti region is a part of the larger Trans-Himalayan drylands across Central Asia (Fig. S1). This high-altitude grazing ecosystem (elevation 4400-4800 m) experiences cold and semiarid climate with temperature dropping below −30 °C in winters. Vegetation growth season starts in May, peaks in July-August, and senescence begins in September. Vegetation consists of grasses (e.g., *Festuca*, *Elymus*, *Stipa*), sedges (*Carex*, *Kobresia*), forbs (*Artemisia*, *Arnebia*, *Eurotia*), and shrubs (*Caragana*, *Rosa*), while trees are rare or absent. Large mammalian herbivores comprise of native species (bharal *Pseudois nayaur*, ibex *Capra sibirica*, and yak *Bos grunniens*) and livestock (goat, sheep, donkey, horse, cattle, and yak-cattle hybrids) (Bagchi and Ritchie, 2010a, 2010b).

After 14 years of the herbivore-exclusion experiment, in 2019, we sampled soil from *n*=10 paired-adjacent plots, in May (early-season), July (peak-season), and September (end-season). We used a 2.5 cm diameter and 20 cm depth corer for soils meant for microbial functions and edaphic variables. We packaged a smaller sample in sterilized vials for DNA analysis.

### Soil microbial functions

We measured soil microbial functions and functional diversity using n = 30 metabolic substrates (Degens and Harris, 1997; Degens, 1998). In this way, we obtained metabolic profiles from 30×10×3×2=1800 measurements, relative to corresponding controls (Fig. 2). We used substrates from different chemical groups: amino acids (Alanine, Arginine, Serine, Asparagine, Glutamine, Glutamate, Histidine), amines (Glucosamine, Putrescine), alcohols (Xylitol, Mannitol, Sorbitol, Glycerol), carbohydrates (D-Glucose, Xylose, Mannose, Sucrose, Inositol), carboxylic acids (α-Ketoglutaric acid, Fumaric acid, Succinic acid, Tartaric acid, Oxalic acid, Malic acid, Ascorbic acid, Citric acid, Malonic acid), an aromatic compound (Inosine), a phenolic compound (Gallic acid), and a polymer (Polysorbate). We measured soil respiration after addition of substrate (substrate induced respiration, SIR) relative to controls (basal respiration) (Degens and Harris, 1997). A substrate was judged to be catabolized if the corresponding SIR was above the 95% confidence-interval of basal respiration (i.e., treatment > control). We used the substrate concentrations such that equal amounts of C was added across all treatments (Degens and Harris, 1997): 100 mM for carboxylic acids and alcohols; 75 mM for carbohydrates; 40 mM for polymers; 15 mM for aromatic, amino acids, amines, and phenols. We pre-incubated 5 g soil at 60% water holding capacity at 28 °C for 24 h. Next, we added 1 ml of substrate solution and incubated soils in air-tight containers along with of 1.5 ml Cresol Red indicator dye (prepared from 2.5 mM NaHCO_3_, 150 mM KCl, 12.5 μg ml^-1^ Cresol Red dye) at 28 °C for 15 h (Rowell, 1995). Next, we estimated substrate utilization (mg C g^-1^ soil day^-1^ respired as CO_2_) from colorimetric measurement of Cresol Red indicator at 570 nm wavelength, alongside corresponding controls and standards (Rowell, 1995). Between these 30 substrates we cover a wide range of metabolic processes that are representative of the microbial repertoire. To ascertain whether our choice of 30 substrates affords robust information on functional diversity we did a thought experiment using iterative resampling of subsets from our data. We randomly sampled 100 combinations of 2 to 29 substrates from the full dataset to generate 2800 subsets of varying compositions (Naidu et al., 2022). For each subset we asked whether microbial species composition is related to functional diversity (see statistical analyses below), to determine how many substrates (and combinations) are necessary for reliable inference.

### Soil DNA extraction, amplification, and sequencing

We estimated microbial taxonomic diversity from operational taxonomic units (OTU) in DNA extracted from soil. We obtained taxonomic profiles from 10×3×2=60 samples, plus corresponding controls. We used MOBIO Power Soil DNA Extraction Kit (Carlsbad, USA) following manufacturer’s instructions. We amplified the 16S rDNA hyper-variable regions V3-V4 using 25 ng DNA extract, KAPA HiFi HotStart Ready Mix, and 100 nM of 341F (5’- CCTACGGGNGGCWGCAG-3’) and 785R (5’-GACTACHVGGGTATCTAATCC-3’) primers (Klindworth et al., 2013). Initial denaturation was at 95 °C for 5 min followed by 25 cycles of 95 °C for 30 s, 55 °C for 45 s and 72 °C for 30 s, with a final extension at 72 °C for 7 min. We sequenced amplicons using Illumina MiSeq platform and processed the data using Mothur (Schloss et al., 2009) to obtain microbial taxonomy data. The raw sequences were aligned to form contigs, and only contigs with length between 300 bp and 532 bp were retained because they cover the targeted error-free sequence of the 16S rDNA region with high confidence. We checked contigs for chimeras and removed them using the UCHIME algorithm (Edgar et al., 2011). We categorized the contigs into OTUs with 97% sequence similarity using the SILVA_v138 database (Quast et al., 2013). As an additional level of quality filtering, we removed the OTUs with cumulative abundance of less than 20 across all samples in the data set. Controls showed minimal absorbance at 260 nm (<1 ng/μl of DNA) compared to soil samples (24 to 70 ng/μl of DNA), and we did not sequence them since they did not show any amplification.

### Edaphic variables

Since edaphic factors can influence variation in microbial communities and in decomposer functions (Fierer and Jackson, 2006; Sinsabaugh et al., 2008), we measured pH, soil moisture, and soil texture, following standard procedures (Robertson et al., 1999). Briefly, we measured soil moisture at monthly intervals as volumetric water content with a time domain reflectometry probe (Spectrum Technologies, USA). We measured pH from 1:2.5 (v/w) aqueous suspension using a table-top pH meter, and soil texture from 50 g soil in 1000 ml of (NaPO_3_)_6_ solution using a Bouyoucos hydrometer (Fig. S2).

### Statistical analyses

We summarized microbial functions with two indices (i) Functional richness: calculated as the number of substrates metabolized. (ii) Dispersion: multivariate measure of functional heterogeneity, estimated as the distance of each sample relative to their respective group centroid (Anderson et al., 2006). We used ‘vegan’ library in R 3.6.3 to estimate richness and dispersion (Oksanen et al., 2019).

We estimated microbial species richness as the number of OTUs in a sample. We used Generalized Linear Models (GLM) with grazing (grazed/ungrazed) and season (early, mid, and late) as fixed-effects, alongside plot-identity as the random-effect to assess the effects of grazing and season on species richness and functional diversity. This GLM accounts for the paired spatial structure in the data (Pinheiro and Bates, 2000), and we used ‘nlme’ library in R 3.6.3. We verified whether the data met the assumptions of GLMs from theoretical and observed quantiles of residuals, and the data did not require any transformations.

### Microbial composition and function

We used ordination with Principal Coordinates Analysis (PCoA) of pair-wise Bray-Curtis dissimilarity to visualise species composition and decomposer functions across seasons and grazing treatment in a multivariate space summarized as two axes (Legendre and Legendre, 2012). We used kriging to visualize the relationship between species composition and microbial functions. Kriging is an interpolation method that helps estimate the value of a variable (e.g., functional richness) over a continuous space of microbial species composition using an existing but smaller set of sampled data points (Dale and Fortin, 2014; Li et al., 2023). We used the first and second axes from PCoA of species composition to interpolate the values of functional indices for unmeasured values of species composition. Kriging is a two-step process: first, a variogram is fitted to determine the spatial (species composition space) covariance structure of the sampled data points; second, interpolate values for the unsampled locations across a given species composition space from the weights derived from the spatial covariance structure of the known data points (Dale and Fortin, 2014; Li et al., 2023). For this, we fitted various variogram models by changing the trend removal parameter (no trend removal, 1^st^ order, 2^nd^ order), model function (exponential, spherical, gaussian), and nugget (fixed, unfixed). First step in fitting the variogram model is trend removal. We did trend removal by fitting a polynomial equation to the predictor values (PCoA axes 1 and 2) using ordinary least squares, and therefore the variograms are computed using the residuals. Here, 1^st^ order trend removal will remove linear trends, whereas 2^nd^ order will remove quadratic trends. Following trend removal, we did model fitting to the empirically determined semivariogram using any of the three model functions where nugget (y-intercept value) is either fixed at zero or unfixed (variable). For each microbial functional index there were 18 different combinations (3 trend removal parameters × 3 model functions × 2 nuggets) to build a variogram model. We selected the final combination for each functional index where the resulting model had the least sum of squares (Table S1). For all the functional indices we selected 2^nd^ order trend removal with exponential function and nugget fixed at zero. We implemented kriging with “geoR” package in R (Ribeiro Jr et al., 2022).

### Distance-based structural equation models

We used distance-based Structural Equation Models (dbSEM) to evaluate whether and how species richness and composition relate to functions, and whether grazers affect the relationship directly or indirectly through edaphic factors. dbSEM is favoured over other types of path analyses since it offers more robust tests of hypothesized causal relationships between variables by using the dissimilarity between samples. dbSEM uses pairwise-dissimilarity matrices for both dependant and predictor variables to disentangle direct and indirect hypothesized causal relationships (Fourtune et al., 2018; Jabot et al., 2020). We used permutation based dbSEM where SEM is performed on the input matrices of variables where rows and columns are randomly permuted to obtain an unbiased P-value for each hypothesized path as the proportion where permuted path coefficient values are less than or equal to the observed path coefficient value (Fourtune et al., 2018; Jabot et al., 2020). The edaphic variables were soil moisture, pH and texture. For soil texture we only included sand content in the analysis because of autocorrelation with clay and silt content. The paths in the dbSEM were motivated by processes that shape microbial communities and were known *a-priori*: (i) Large mammalian herbivores alter microbial communities (Grazing → Species diversity (species richness, species composition)) (Sankaran and Augustine, 2004; Bagchi et al., 2017). (ii) Herbivores can regulate decomposition process directly (Grazing → Function (functional richness, functional composition)), or indirectly through species composition (Grazing → Species diversity → Function) (Bagchi et al., 2017; Naidu et al., 2022; Roy and Bagchi, 2022). (iii) Background edaphic variables can influence microbial communities (Edaphic [pH, sand, moisture] → Species diversity) (Girvan et al., 2003; Fierer and Jackson, 2006; Bagchi et al., 2017). (iv) Variation in background edaphic variables can alter function directly (Edaphic → Function), or indirectly through influence on species diversity (Edaphic → Species diversity → Function) (Girvan et al., 2003; Delgado-Baquerizo et al., 2016b; Bagchi et al., 2017). (v) both microbial community and functions can vary temporally (Season → Species diversity, Season → Function) (Bardgett et al., 2005; Frossard et al., 2012; Delgado-Baquerizo et al., 2013; Bagchi et al., 2017). (vi) Grazing can also influence soil moisture (Grazing → Moisture) (Bagchi et al., 2017), that can in-turn influence pH (Moisture → pH). (vii) Sand content can also alter moisture (Sand → Moisture) and pH (Sand → pH). We assessed goodness of fit between the model and data with χ^2^ test, comparative fit index (CFI), and Standardized Root Mean Square Error of Approximation (RMSEA). We accepted the model P(χ^2^)> 0.05, CFI ∼ 1, and RMSEA < 0.05. We judged the agreement between data and individual hypothesized paths at α=0.1 to account for high variability in the individual dissimilarity matrices. We implemented paths for dbSEM using ‘lavaan’ library in R (Rosseel, 2012).

As a thought experiment we evaluated whether our dataset of 30 functions was adequate to understand if species richness and/or composition affected functions, or not. We first resampled our microbial function data to obtain datasets with different number of functions (from 2 functions to 29 functions, i.e., 28 subsets). Since each subset can be envisioned as a study covering a certain number of functions, we evaluated 100 different possible combinations thereof (Naidu et al., 2022). So, for each subset, we randomly iterated 100 different combinations of functions, that ultimately generated 2800 datasets (28 subsets × 100 combinations of each). We then repeated dbSEM analysis for each subset to ask whether the number of functions can influence interpretation of diversity-function relationships in each of these 2800 datasets. Specifically, after each dbSEM iteration of a subset, we checked whether the path connecting species diversity (species richness, and species composition) to functions (function richness, and functional composition) was significant, or not (α=0.1). Since, previous evidence for a positive relationship between species composition and functions come from metagenomic extrapolations of 10^3^ predicted functions, this thought experiment allows us to evaluate whether the relationship holds true for 10^1^ actual functions (and various combinations), or not (Fierer et al., 2013; Galand et al., 2018).

## Results

### Microbial communities and functions *–*

We recorded 1937 unique OTUs across 47 known phyla and 7650894 reads. Of these, only 924 OTUs were identified to genus level. Actinobacteriota, Acidobacteriota, Planctomycetota, Proteobacteria and Verrucomicrobiota were most common (Fig. 1, Fig. S3). Species richness of OTUs ranged between 494 and 866 per sample (mean 713±80 SD). Similarly, functional richness (i.e., number of substrates metabolized) ranged between 6 and 30 (mean 21±7 SD, Fig. 1, Fig. S3). Expectedly, microbial activity as SIR was high for carboxylic acids, and low for amino acids (Fig. 1, Fig. S3).

**Figure 1.**
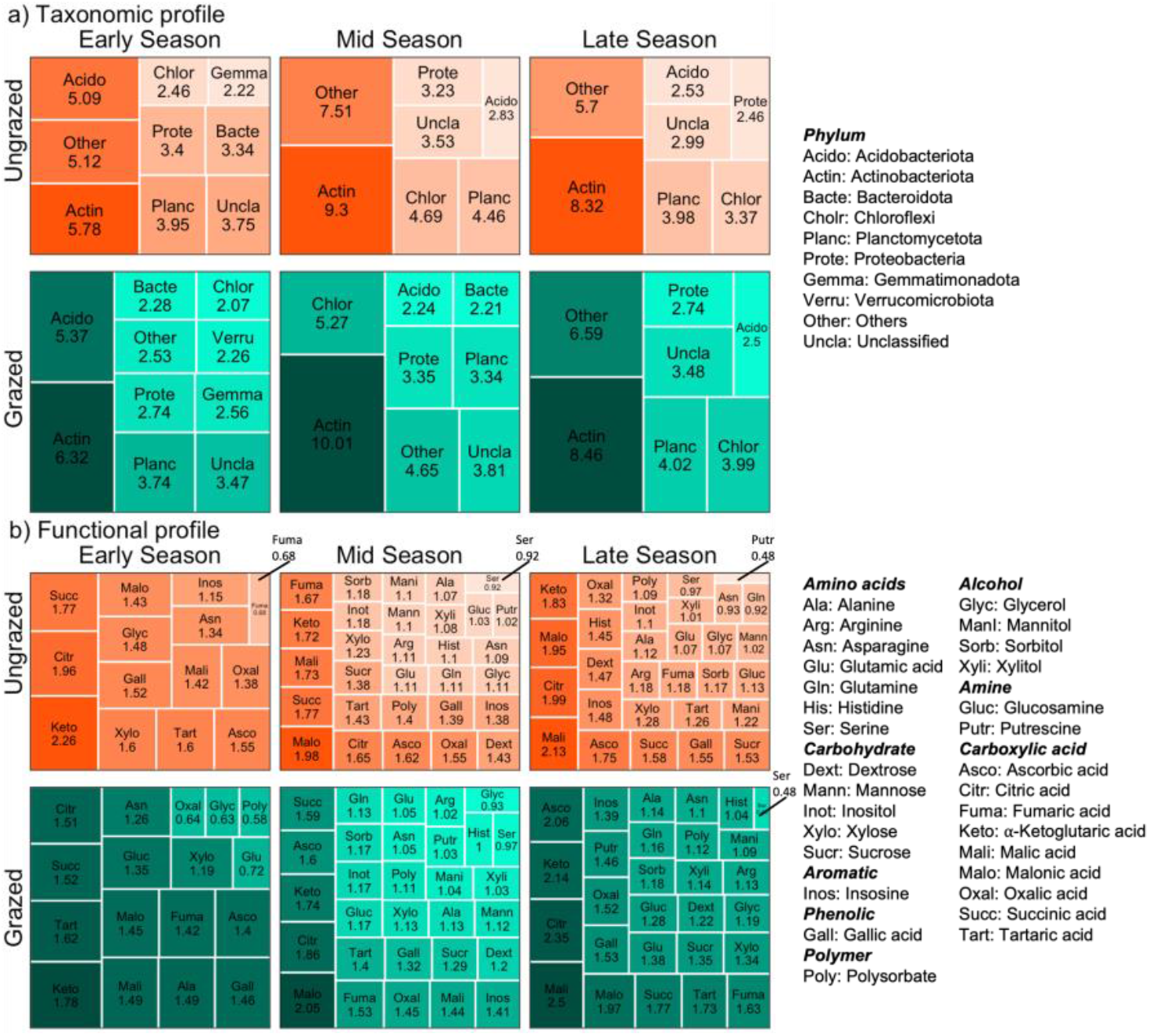
Treemaps summarizing the microbial taxonomic profile (phylum level) and microbial functional profile (n=30 substrates) in the presence and absence of large mammalian herbivores averaged for all plots and across the growth season (Early, Mid, and Late). Area of each cell is proportional to the total abundance of each phylum and catabolic activity for each substrate, respectively. Number denotes the percentage of the relative abundance (taxonomy) and relative microbial activity (function). Data are from n=10 paired grazed-and-fenced plots in Spiti region of Trans-Himalaya in northern India.

### Seasonality and herbivore-effects on microbial species and functions *–*

Variation in species (OTUs) richness was neither explained by season (Fig. 2a; F_2,45_=0.02, *P*=0.98), nor by grazing (Fig. 2a; F_1,45_=1.61, *P*=0.21). Microbial functional richness varied with season (Fig. 2b; F_2,45_=38.96, *P*<0.001) but did not vary with grazing (Fig. 2b; F_1,45_=0.01, *P*=0.98). Similarly, microbial functional dispersion varied with season (Fig. 2c; F_2,45_=48.89, *P*<0.001) but did not vary with grazing (Fig. 2c; F_1,45_=0.01, *P*=0.92). In multivariate space summarized with PCoA ordination, microbial species composition varied with season (Fig. 3; F_2,54_=4.70, *P*=0.001), but not with grazing (Fig. 3; F_1,54_=1.33, *P*=0.21). Functional composition also varied with season (Fig. 3; F_2,54_=4.83, *P*=0.001), but not with grazing (Fig. 3; F_1,54_=0.87, *P*=0.55).

**Figure 2.**
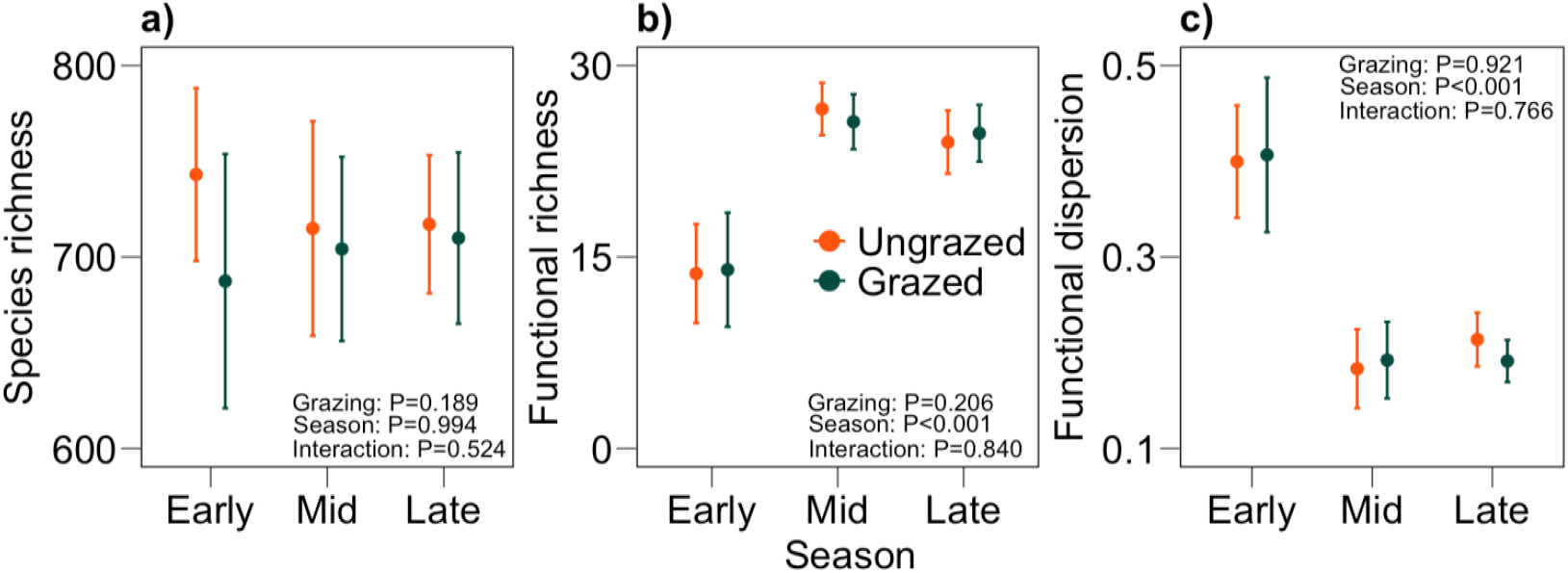
Species richness of soil microbes (a), functional richness (b), and functional dispersion (c), in the presence and absence of large mammalian herbivores across the growth season. Species richness (mean ± 95% CI) did not vary between grazed and ungrazed plots, and with the growth season. Functional richness and functional dispersion did not vary between grazed and ungrazed plots, but varied across the growth season. Data are from n=10 paired grazed-and-fenced plots in Spiti region of Trans-Himalaya in northern India.

**Figure 3.**
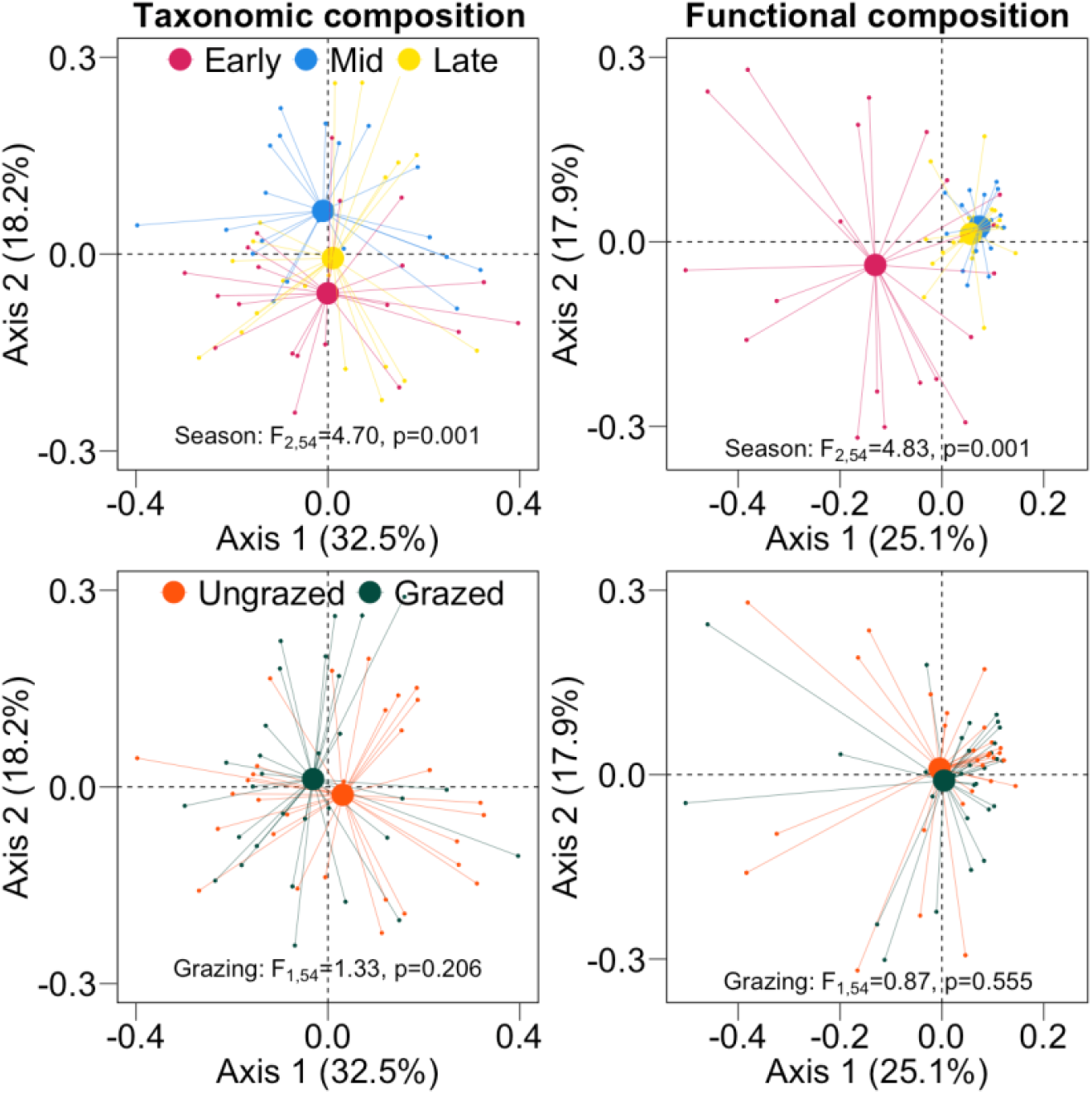
Summary of ordination by Principal Coordinates Analysis (PCoA) for microbial communities and decomposer functions in the presence and absence of large mammalian herbivores, and at different points in the growth season. Overall, both taxonomic and functional composition of soil microbial decomposers did not vary between grazed and ungrazed plots, but they varied across the growth season. Data are from n=10 paired grazed-and-fenced plots in Spiti region of Trans-Himalaya in northern India.

### Relationship between species composition and functions *–*

Kriged maps of microbial functions indicated that differences in community composition were related to differences in functions (Fig. 4), and these were further assessed using dbSEM (Fig. 5). Overall, dbSEM for functional richness, and functional dispersion (Fig. 5) indicated good match between the data and hypothesized paths (SRMR=0.005, RMSEA<0.001, P-value (RMSEA)=0.982, CFI>0.999, χ^2^=1.34, P-value (χ^2^)=0.512). Path coefficients suggested a positive relationship between species composition and functional richness (Species composition → Functional richness; path coefficient=0.08, *P*=0.06; Fig. 5), but not with functional composition (Species composition → Functional composition; *P*>0.10; Fig. 5). Species richness did not have any direct or indirect effect on either functional richness or functional composition (*P*>0.10; Fig. 5). Soil texture had a positive effect on species composition (Texture → Species composition; path coefficient=0.09, *P*=0.001; Fig. 5). Seasonality had positive relationship with species composition (Season → Species composition; path coefficient=0.14, *P*<0.001; Fig. 5), functional composition (Season → Functional composition; path coefficient=0.25, *P*<0.001; Fig. 5), and soil moisture (Season → Moisture; path coefficient=0.29, *P*<0.001; Fig. 5), such that as the season progresses from early- to mid- to late-season, species composition, functional composition, and soil moisture becomes more dissimilar. But seasonality had a negative relationship with species richness (Season → Species richness; path coefficient=-0.04, *P*=0.08; Fig. 5), such that as the season progresses from early- to mid- to late-season, species richness between samples becomes more similar. Variation in moisture influenced both the species composition (Moisture → Species composition; path coefficient=0.12, *P*=0.02; Fig. 5), and species richness (Moisture → Species richness; path coefficient=0.08, *P*=0.09; Fig. 5). In this way seasonality had a direct (Season → Species composition → Function), as well as indirect effect on the diversity-function relationship through moisture (Season → Moisture → Species composition → Function). Variation in pH had a negative relationship with Species richness (pH → Species richness; path coefficient=-0.07, *P*=0.09; Fig. 5). Grazing did not influence this relationship directly (Grazing → Species composition → Function), or indirectly through edaphic factors (Grazing → Edaphic → Species composition → Function).

**Figure 4.**
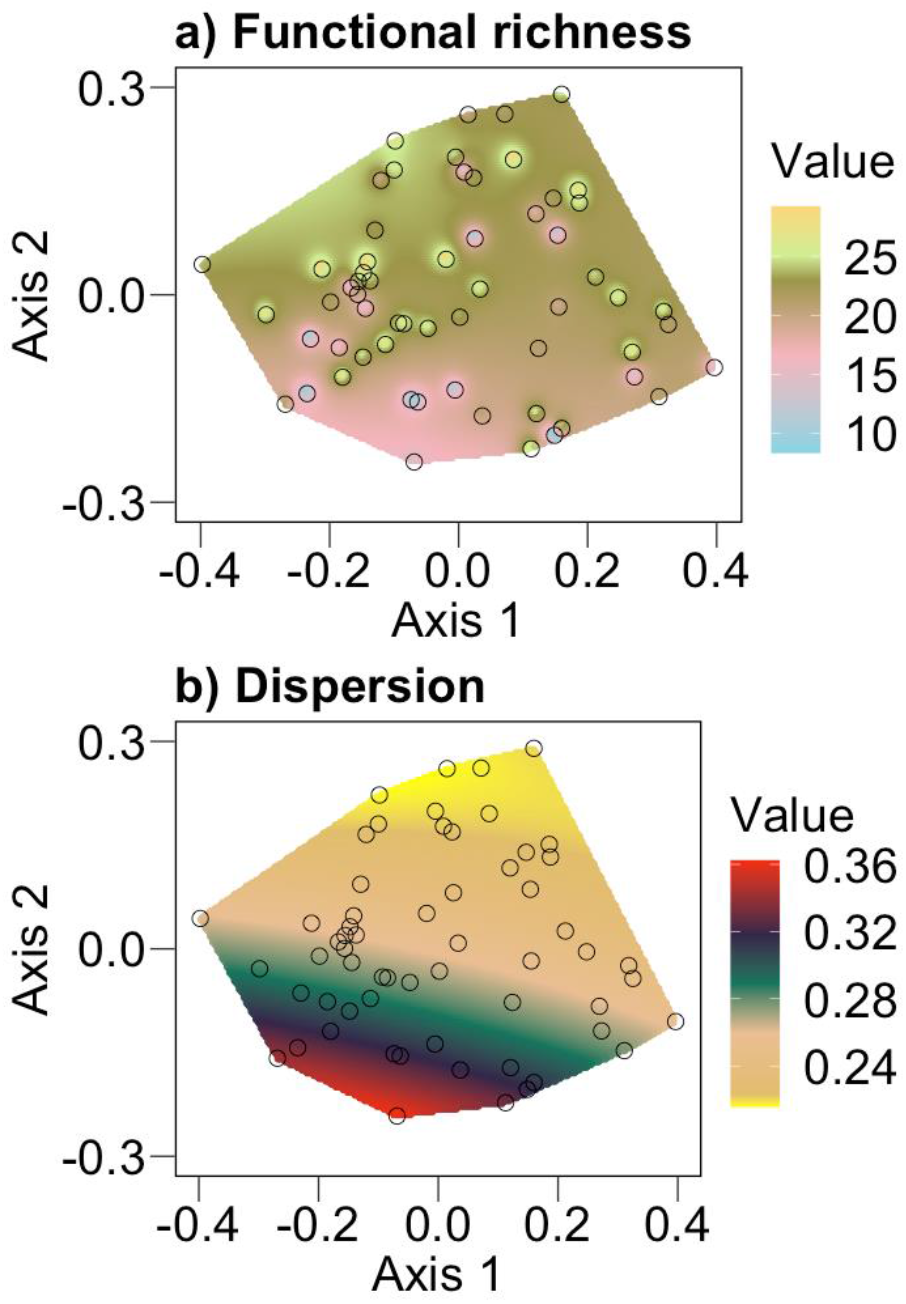
Kriged maps summarizing the relationship of microbial functional richness (a), and functional dispersion (b) with microbial species composition, show that microbial decomposer functions can vary with species composition. The colored space highlights the interpolation of microbial functions across the extent of microbial species composition from the study site measured at different points in the growth season. Species composition is represented as first and second axes of principal coordinate analysis (PCoA). Circles denote the sampled data points for species composition. Value in the legend indicates the range of interpolated values of respective functional index.

**Figure 5.**
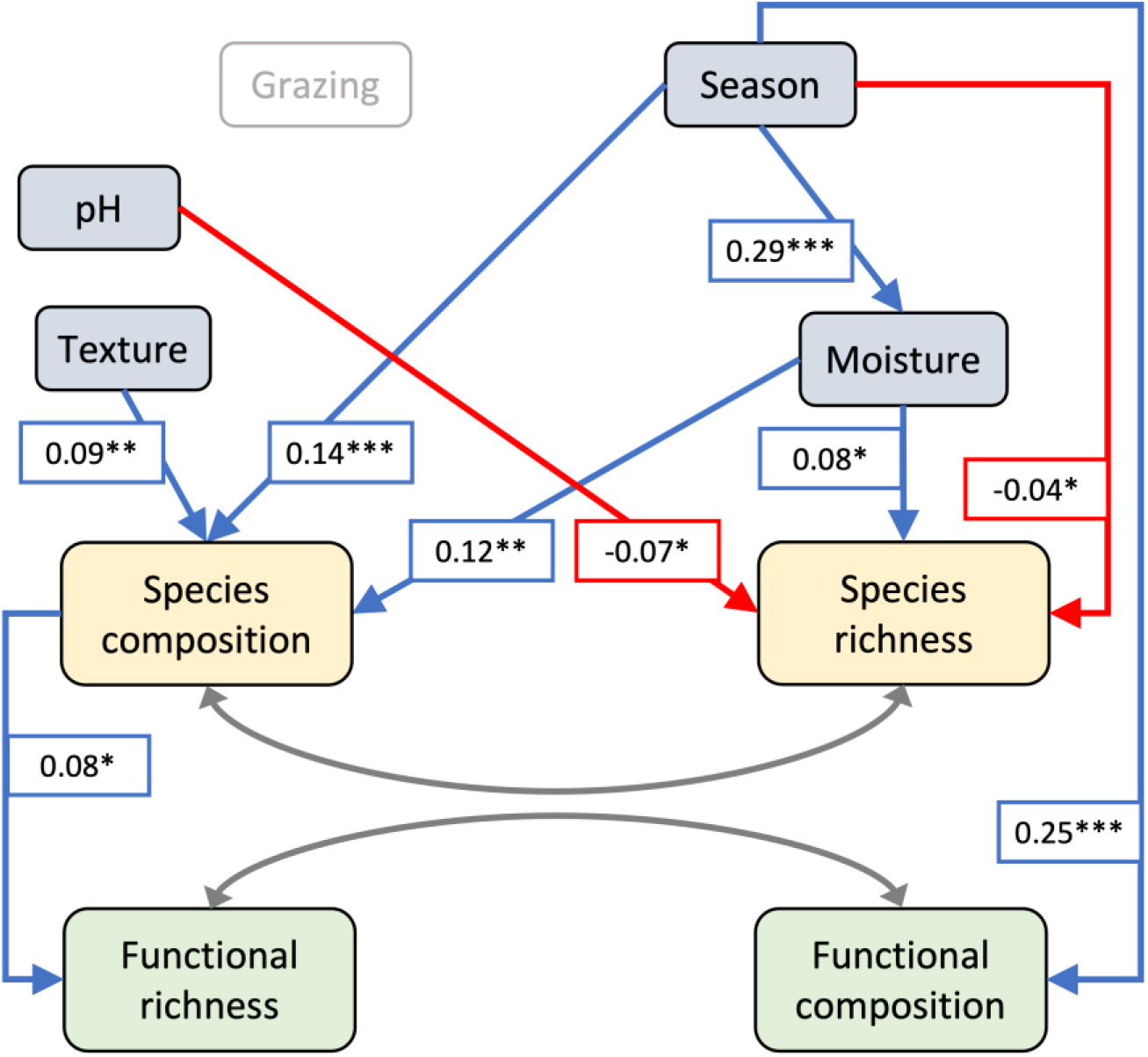
Summary of distance-based structural equation model (dbSEM) to evaluate the influence of grazing, seasons, soil abiotic properties, on the relationship between species diversity and microbial functions. Values represent standardized path coefficients and asterix represent statistical significance (***P≤0.01, **P≤0.05, and *P≤0.10). Modelled paths from grazing (grazed/ungrazed) were non-significant, and are not shown. Diagnostics for the model was SRMR=0.004, RMSEA<0.001, P-value (RMSEA)=0.982, CFI>0.999, χ^2^=1.34, df=2, P-value (χ^2^)=0.512. Overall, there was a positive relationship between microbial functions and species composition, whereas species richness had no influence on functions.

Through our thought experiment involving iterative resampling of the functional data, we find that 16 or more substrates in different combinations can provide robust interpretations of functional diversity (Fig. 6). The relationship between species composition and functions (Fig. 5) could be inferred as long as there is quantitative information from 16 or more functions (Fig. 6).

**Figure 6.**
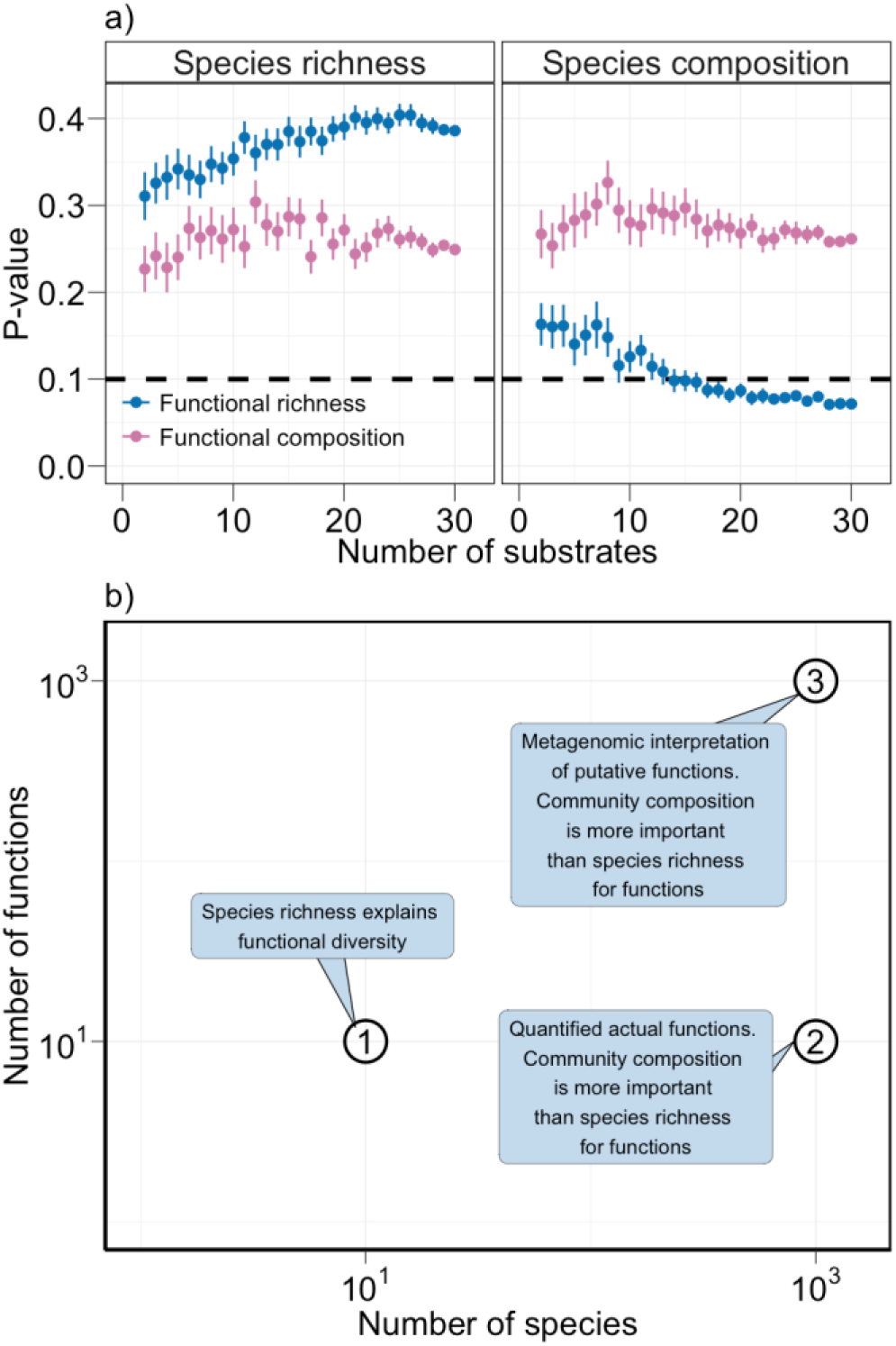
Summary of thought experiment to assess how the number of measured functions can influence the interpretation of diversity-function relationships in (a). Through iterative resampling of our functional data, we generated 2800 subsets with 100 different possible combinations of 2 to 29 functions. For each subset, we repeated dbSEM analyses (Fig. 5) checked for the significance (α=0.1) of path connecting species diversity (species richness, and species composition) to functions (function richness, and functional composition). We report the P-value of the above-mentioned paths (Species diversity → Function) for each combination of 2 to 29 functions (mean ± 95 CI across 100 iterations). This thought experiment reveals that 16 or more functions are sufficient to recapitulate the results seen for the full dataset of 30 substrates. Conceptual summary of three types of studies on diversity-function relationships for microbes that accommodate different number of species and functions (b). Studies on experimentally assembled communities of 10^1^ species and 10^1^ functions (Delgado-Baquerizo et al., 2016a; Wagg et al., 2019), usually encounter a positive relationship between species richness and functions (i.e., #1). Studies with hyper-diverse natural communities with 10^3^ species (Delgado-Baquerizo et al., 2017; this study), generally encounter partial redundancy (i.e., #2). Studies with metagenomic interpretations of a large number of putative functions (Fierer et al., 2013; Louca et al., 2016; Galand et al., 2018; Chen et al., 2022) also encounter partial redundancy (i.e., #3).

## Discussion

While diversity-function relationships in producers, and their modulation by consumers at higher trophic levels, are well known, they remain uncertain for microbial decomposers and decomposition functions. Here we investigate whether, or not, microbial taxonomic diversity is related to microbial decomposition functions and find three key results: (i) soil microbial communities and functions are more variable across time than over space (Fig. 1 – 3). (ii) microbial species composition affects functional richness, but not functional composition (Fig. 4 – 5, Fig. S4). And, (iii) unlike background edaphic variables and seasonality, large mammalian herbivores do not influence the diversity-function relationship for microbes (Fig. 5). Additionally, a small subset of functions (16 or more substrates) covering an array of decomposer metabolic processes provide a sufficient sample-size to understand diversity-function relationships in soil microbes (Fig. 6).

We find temporal variation in microbial community structure and functions to be more pronounced than spatial heterogeneity (Figs. 2, 3, 5). This is consistent with other studies where fluctuations in resource availability explained why microbes respond to temporal variables more than spatial variables (Frossard et al., 2012; Bagchi et al., 2017; Roy and Bagchi, 2022; Roy et al., 2023). Further, we find a consistent effect of soil moisture and texture on the decomposer diversity-function relationship (Fig. 5). Moisture and texture can play an important role in habitat-filtering (Bass-Becking, 1934) as they promote functions through taxonomic composition (Fig. 5), resulting in the loss of functional redundancies among decomposers. While the effect of moisture and texture (Fig. 5) is consistent with other studies across ecosystems (Girvan et al., 2003; Delgado-Baquerizo et al., 2016c), unlike other studies (Fierer and Jackson, 2006; Sinsabaugh et al., 2008) we find no effect of soil pH on the relationship. This could be due to relative invariance in soil pH across the sampled plots (space), but it can vary more substantially between different landscapes and ecosystems.

The lack of spatial heterogeneity in hyper-diverse microbial community structure and functions can result in functional redundancy. As a case in point, we recorded more than 700 OTUs on average per sample, and 1937 OTUs overall across 7.6 million reads. Therefore, and understandably, the data did not show any support for the relationship between species richness and functions (Fig. S4). For instance, where local richness is below 200 OTUs per sample, there is evidence of coupling and therefore a lack of redundancy (Delgado-Baquerizo et al., 2016a). But, further along at 700 OTUs per sample (Fig. 3 – 4) microbial diversity can appear de-coupled from functions (Peter et al., 2011; Louca et al., 2016; Chen et al., 2022). Thus, not surprisingly, different studies have encountered coupled as well as de-coupled relationships between microbial species richness and functions (Peter et al., 2011; Delgado-Baquerizo et al., 2016a; Louca et al., 2016; Chen et al., 2022), and our data emphasize how it is important to accommodate species composition (Hooper et al., 2005).

The existence of redundancy has often been challenged for hyper-diverse natural microbial communities (Fierer et al., 2013; Galand et al., 2018). In its stringent form, functional redundancy states that in a community there exist organisms who perform the same set of functions and can be easily replaced without any changes in the function (Allison and Martiny, 2008; Galand et al., 2018). However, this is seldom the case in natural communities since the environmental thresholds (such as temperature, precipitation) of different organisms to perform the same set of functions are different. For instance, we find that seasonality and edaphic factors (moisture and texture) exert strong control over the diversity-function relationship. Moreover, the notion of functional redundancy is also dependent on which functions were investigated, and how many (Fig. 6). This is because in natural communities, microbes can perform multiple functions at once where in one hand, some of the functions would be shared by other microbes making the function redundant. On the other hand, some of the functions would be specific to only a select few making the function specialized and non-redundant. Therefore, instead of complete redundancy our results suggest partial redundancy. But, since large number of the 1937 OTUs have not yet been resolved to genus/species level, we do not fully know the identity of microbes that are responsible for higher functions. However, anticipated technological advances can bridge this knowledge gap and can foster better interpretations of the relationship (Thompson et al., 2017).

Given the low resolution of microbial taxonomy is soil, previous studies have often relied on metagenomic interpretations and extrapolations of putative functions from the presence/absence of genetic code for various proteins and enzymes (Fierer et al., 2013; Galand et al., 2018). This does not automatically guarantee that the corresponding functions are actually performed in a sample because ‘everything is everywhere, but environment selects’ (Bass-Becking, 1934). We assessed the diversity-function relationship by measuring the magnitude of functions, albeit for a small number of 30 functions. But our thought experiment suggests that interpretations of the relationship is generalizable and reliable for 10^1^ functions compared against 10^3^ putative functions from the metagenomic approach (Fierer et al., 2013; Delgado-Baquerizo et al., 2017; Galand et al., 2018). Further, such coupled diversity-function relationships for microbes and decomposition agrees with patterns seen with producers and production where identity of the species is more important than the number of species (Hooper and Vitousek, 1997; Tilman et al., 1997).

We find that the relationship between microbial diversity and function is robust to the presence or absence of large mammalian herbivores. This also is in line with resilience and resistance patterns among producers and their production where herbivore-exclusion does not impact producer diversity and production (Kohli et al., 2019). However, contrary to the direct effect of consumers on producers, the mechanisms operating for decomposers might be different (Bagchi et al., 2017; Naidu et al., 2022; Roy and Bagchi, 2022). There could be multiple interdependent paths that support such reliance and resistance. First, a balance between the relative strengths of direct and indirect influence of herbivores can manifest as no net effect on the decomposer diversity-function relationship (Bagchi and Ritchie, 2010a; Cherif and Loreau, 2013; Naidu et al., 2022; Roy and Bagchi, 2022; Roy et al., 2023). Second, the tight coupling between microbial diversity and function could have established before the start of our decadal-scale herbivore-exclusion experiment due to competition-cooperation interactions (Morrissey et al., 2019; Lechón-Alonso et al., 2021). Once established, the interactions among microbial decomposers may withstand the experimental exclusion of herbivores (resistance), and/or recover quickly from it (resilience) (Allison and Martiny, 2008; Morrissey et al., 2019; López Zieher et al., 2020). Third, herbivore effects might be different on other types of decomposers which were not quantified by our study, e.g., archaea, uni- and multi-cellular eukaryotic decomposers (Bagchi et al., 2017; Roy et al., 2023).

Overall, we find that differences in microbial community composition affect microbial functional diversity, whereas microbial species richness alone has no effect (Galand et al., 2018). This is inconsistent with a key aspect of functional redundancy that posits no relationship between species diversity (both richness and composition) and functional diversity (Hooper and Vitousek, 1997; Allison and Martiny, 2008; Galand et al., 2018). Instead, our results suggest that loss or gain of some species (i.e., composition) in the microbial community can have larger impacts on microbial functions than others. Therefore, the potential for species diversity to buffer (i.e., resistance) and ameliorate (i.e., resilience) the effect of stressors is contingent upon the identities of the species that remain unaffected by the stressor (Hooper and Vitousek, 1997; Tilman et al., 1997). While microbial diversity-function relationships appear resistant and/or resilient to the experimental removal of large mammalian herbivores, they are sensitive to variability in soil moisture which is a major determinant of most dryland ecosystem processes and services (Allison and Martiny, 2008; Millard and Singh, 2010; Delgado-Baquerizo et al., 2013; Griffiths and Philippot, 2013; Maestre et al., 2016). Future studies can address how resistance and resilience of microbial functions dovetail with various aspects of global change including rising inter-annual variability in precipitation and its intra-annual redistribution. Future changes in soil moisture availability through continued alteration of precipitation in drylands (Schlaepfer et al., 2017; Konapala et al., 2020), particularly in the Trans-Himalaya and Central Asia’s high-elevation drylands, might challenge their conservation and sustainable management for ecosystem functions and services.

## Acknowledgements

SR was supported by a graduate fellowship from CSIR-India in IISc, and Discipline Hopping for Discovery Science grant by NERC-UK in UEA; SB was supported by MHRD-India. Funding support for fieldwork and lab-analyses were obtained from DST-SERB, DST-FIST, DBT-IISc. Govt. of Himachal Pradesh and state departments have supported our research in Spiti. We received generous support from Dorje Chhering, Dorje Chhewang, and Deepak Shetti, in field and in lab.

## APPENDIX

**Figure S1.**
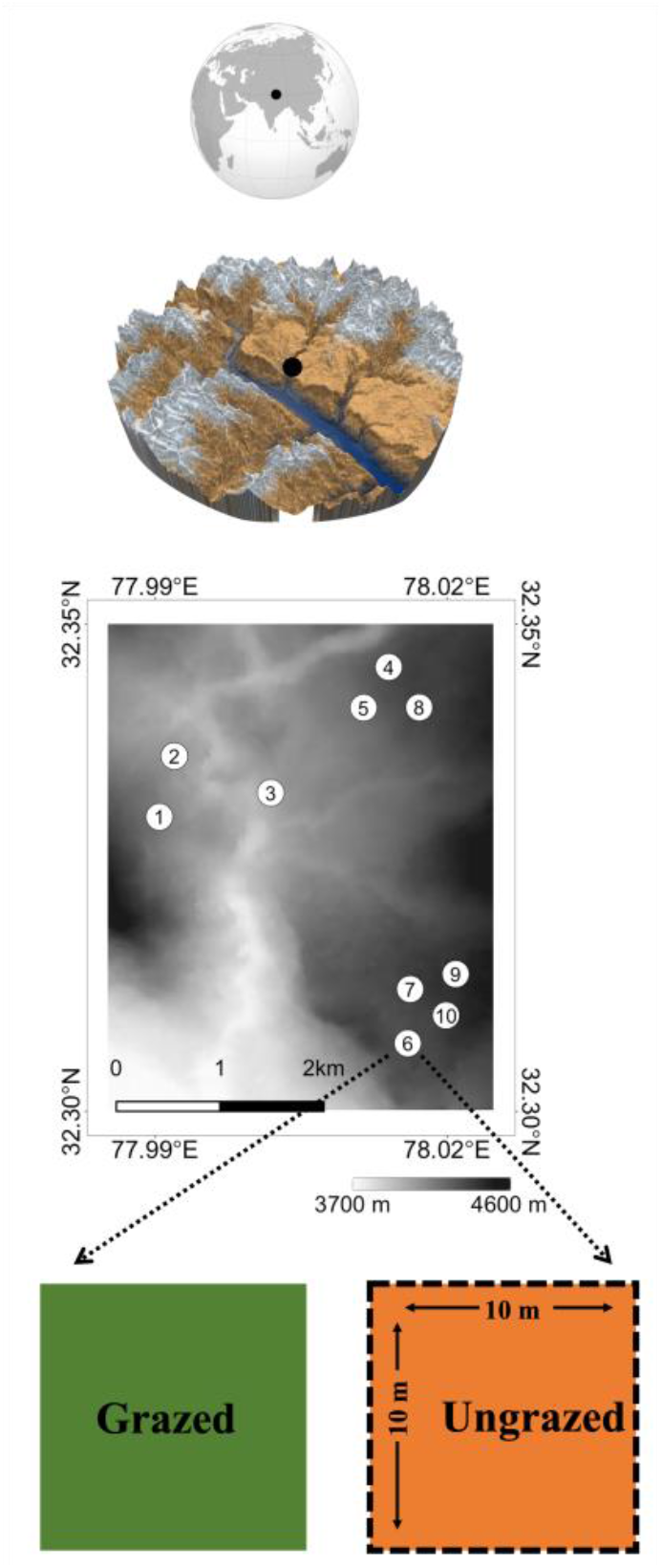
Map of study site showing sampling locations in Spiti region of the Trans-Himalaya in northern India. Each sample consists of a paired grazed-and fenced plot (10 m × 10 m).

**Figure S2.**
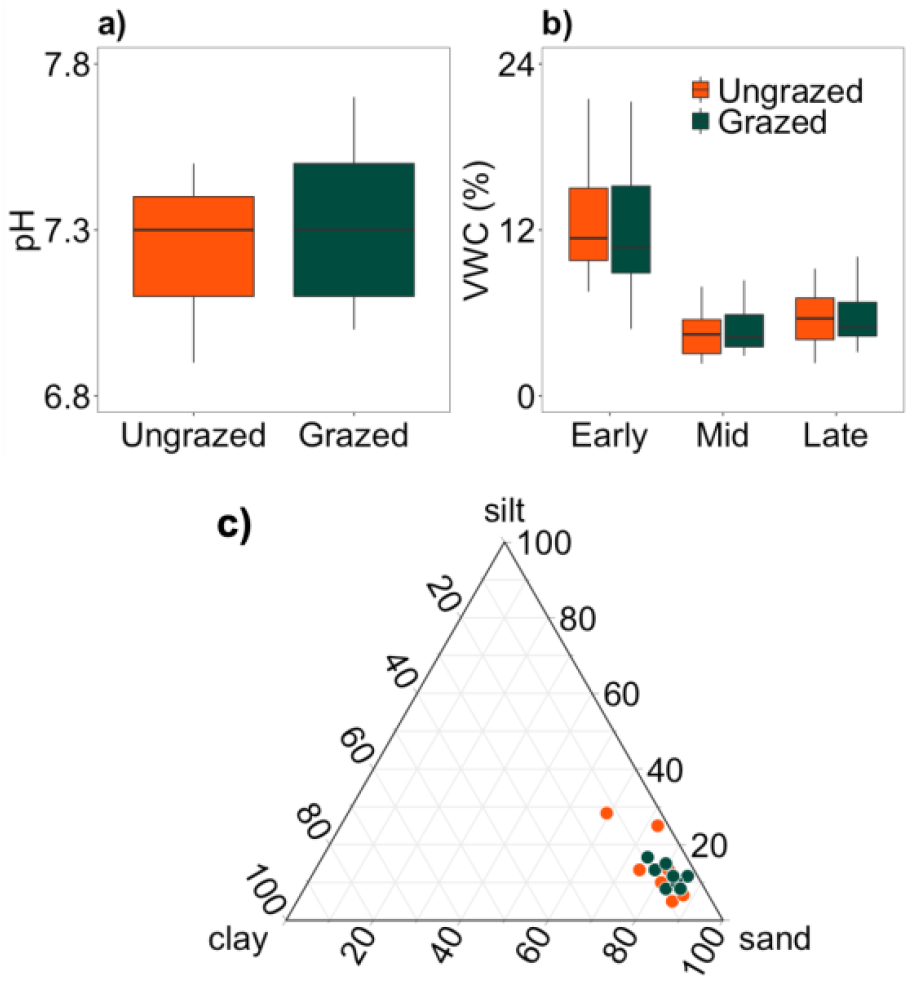
Summary of edaphic conditions – soil pH (a), soil moisture (b), and soil texture (c), in the presence and absence of large mammalian herbivores (n=10 paired plots). Soils were near-neutral, showed higher moisture content during early-season (snowmelt), and were of sandy-loam texture.

**Figure S3.**
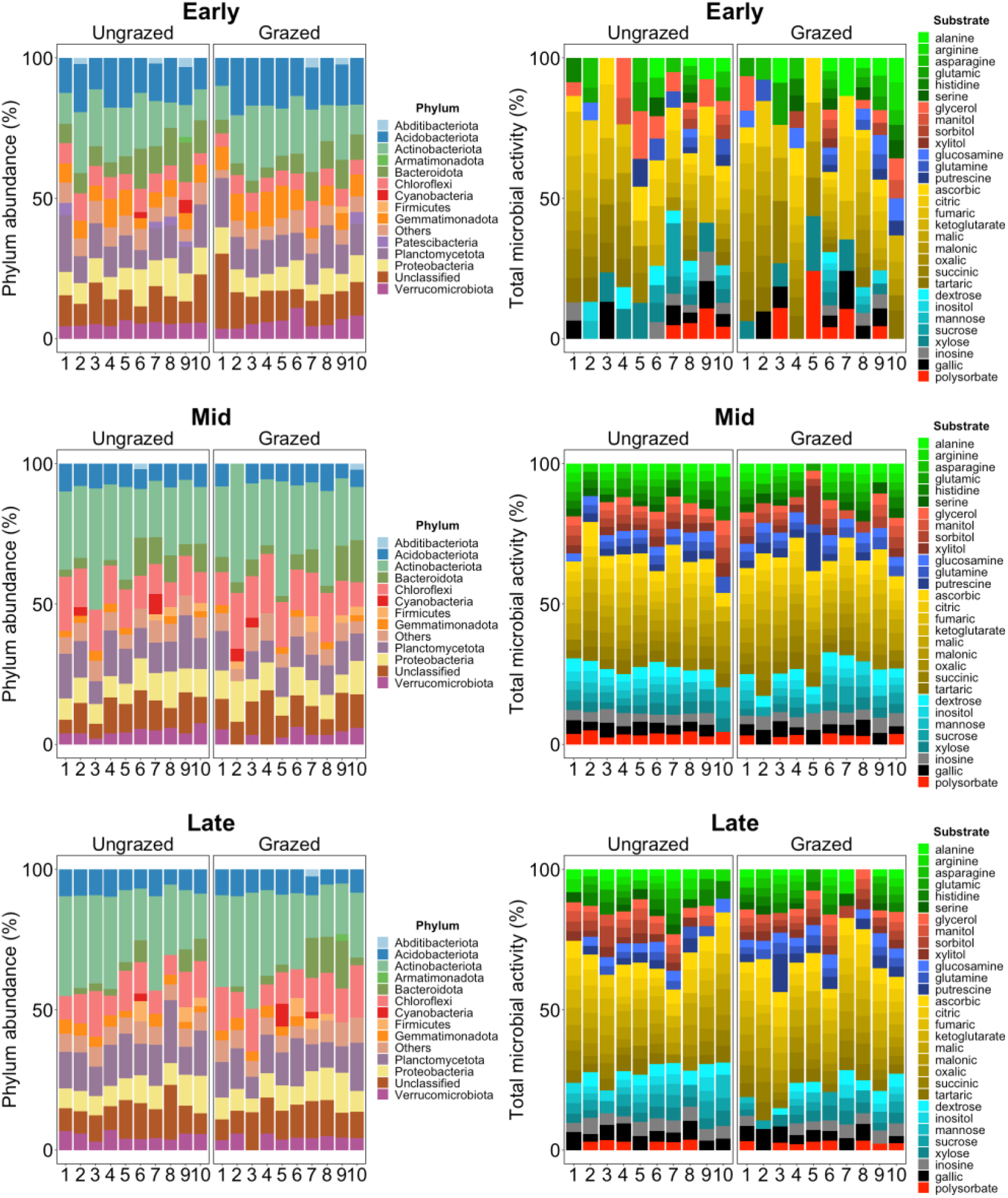
Summary of soil microbial communities at the phylum level (covering >=90% of the total abundance) and microbial functional profiles (n=30 substrates) in the presence and absence of large mammalian herbivores (n=10 paired plots) across the growth season.

**Figure S4.**
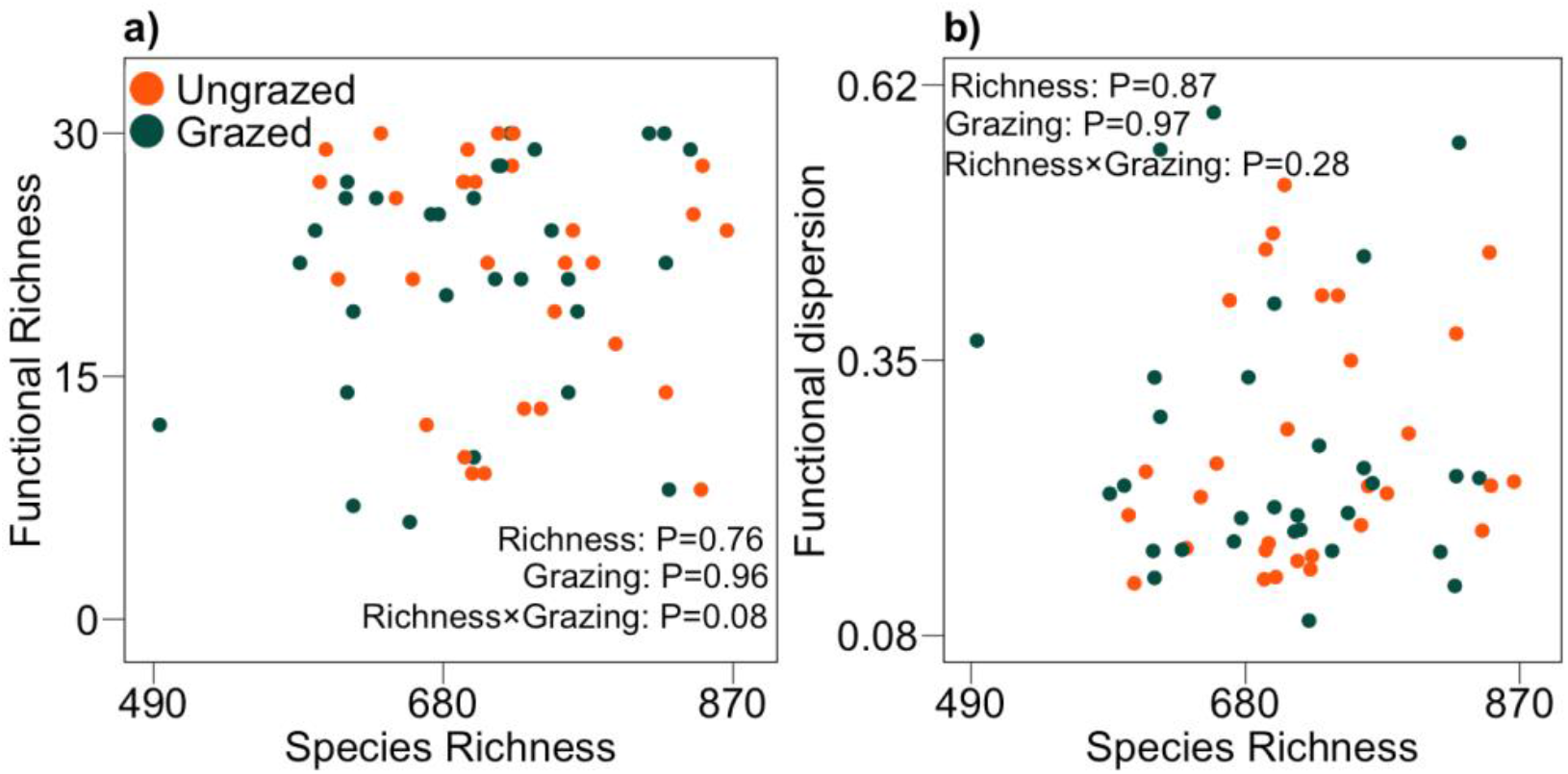
Relationship between species richness and functional richness (a), and functional dispersion (b), in the presence and absence of large mammalian herbivores across the growth season. Species richness was not related to microbial functions in both grazed and ungrazed plots across the growth season.

**Table S1.**
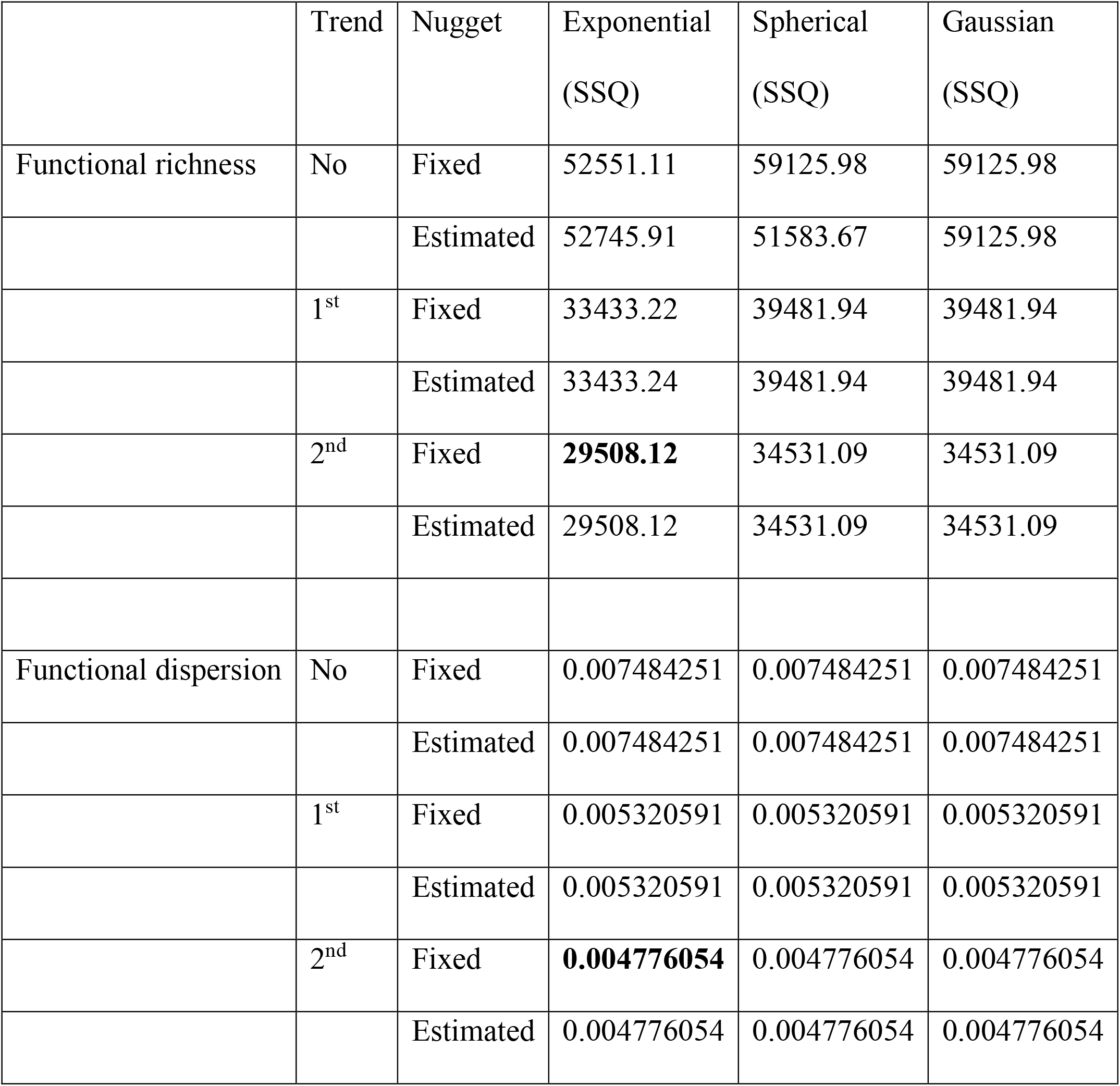
Best model for kriging using sum of squares (SSQ)

**Table S2.**
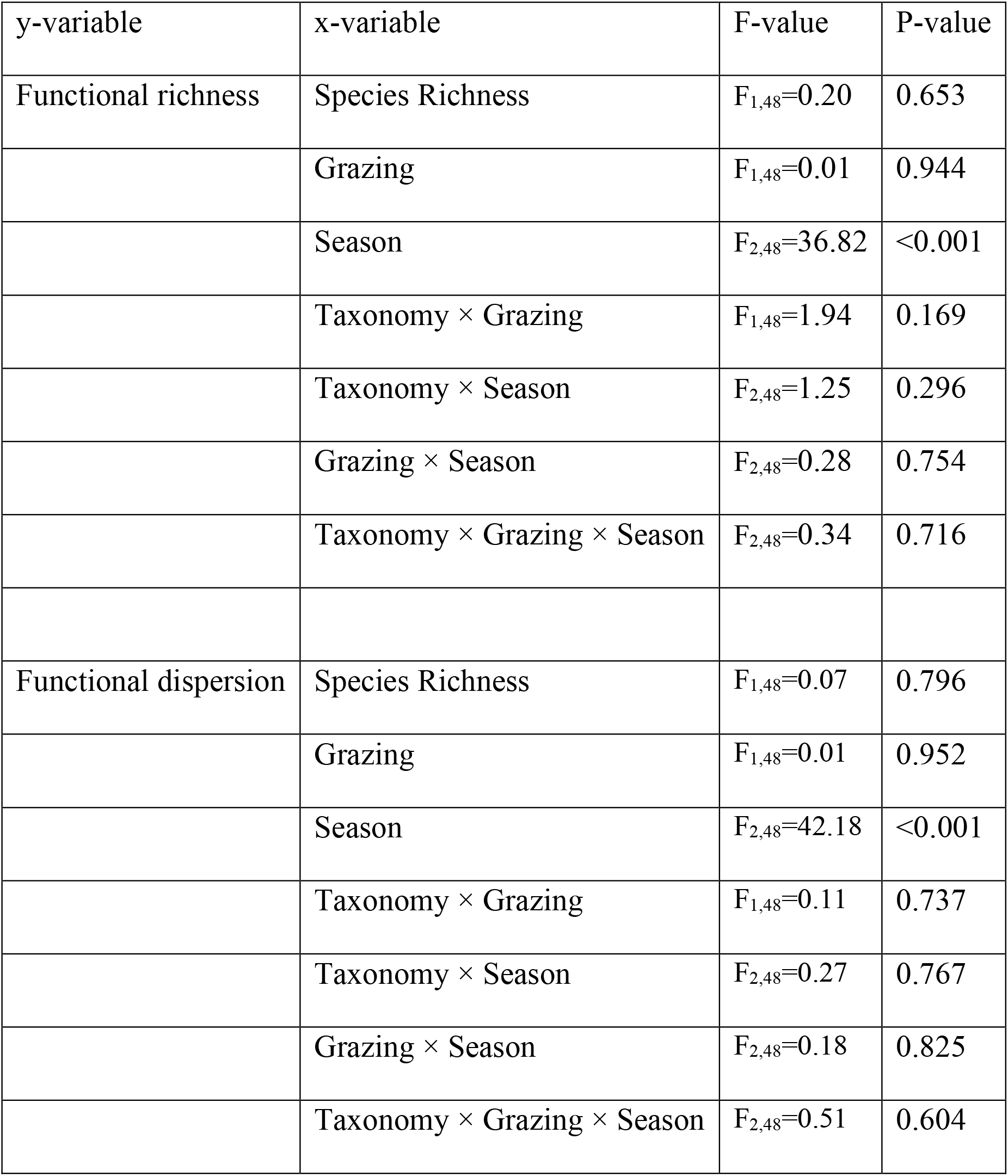
Summary of omnibus GLM model: Functions ∼ Species richness * Grazing * Season

| y-variable | x-variable | F-value | P-value |
| --- | --- | --- | --- |
| Functional richness | Species Richness | $F_{1,48}=0.20$ | 0.653 |
| | Grazing | $F_{1,48}=0.01$ | 0.944 |
| | Season | $F_{2,48}=36.82$ | <0.001 |
| | Taxonomy × Grazing | $F_{1,48}=1.94$ | 0.169 |
| | Taxonomy × Season | $F_{2,48}=1.25$ | 0.296 |
| | Grazing × Season | $F_{2,48}=0.28$ | 0.754 |
| | Taxonomy × Grazing × Season | $F_{2,48}=0.34$ | 0.716 |
| Functional dispersion | Species Richness | $F_{1,48}=0.07$ | 0.796 |
| | Grazing | $F_{1,48}=0.01$ | 0.952 |
| | Season | $F_{2,48}=42.18$ | <0.001 |
| | Taxonomy × Grazing | $F_{1,48}=0.11$ | 0.737 |
| | Taxonomy × Season | $F_{2,48}=0.27$ | 0.767 |
| | Grazing × Season | $F_{2,48}=0.18$ | 0.825 |
| | Taxonomy × Grazing × Season | $F_{2,48}=0.51$ | 0.604 |

